# Characterizing postoperative T and B cell dysfunction in cancer surgery patients, using COVID-19 as a model antigen

**DOI:** 10.1101/2024.12.23.629453

**Authors:** Oladunni Olanubi, Taylor Dion, Regan Macdonald, Rafeah Alam, Christiano Tanese De Souza, Richard Hu, Angela Crawley, Rebecca C Auer

## Abstract

For most cancers, surgery is an effective intervention method for cure but, despite the benefits, many patients recur with metastatic disease. Surgery has been shown to impair immune function by causing suppression of immune cells. In this study, we used a COVID-19 immunological toolkit to answer questions regarding the effects of surgery on antigen specific CD8^+^ T cells and B cells by exploiting the responses to the spike protein in vaccinated cancer patients. We demonstrate that surgical stress results in a reduction in the number of CD8^+^ T cell that produce cytokines and B cells that secrete antibodies in response to antigen. This study will improve our understanding of surgery-induced T cell and B cell dysfunction

## Introduction

Surgery is essential for the treatment of solid malignancies, however, the stress which results from resecting the tumor has unintended consequences on immune function. Our group and others have shown that the suppression of cell mediated anti-tumor immunity observed in the aftermath of cancer surgery is associated with an increased risk of cancer recurrence and metastasis in preclinical models^1-4^. While many of these studies have focused on the effects of surgery on NK cells of the innate immune system, the adaptive and humoral immune system in particular CD8+ T cells and B cells haven’t received much attention. The widespread vaccination against COVID-19 and the availability of immunological assays to measure SARS-CoV2 specific immune responses presents and ideal opportunity to evaluate the effect of surgery on antigen specific CD8^+^ T cell and B cell responses.

## Methods

A cohort of patients undergoing major abdominal surgery for different types of malignancies and had received at least 2 doses of the Pfizer BNT vaccine were included in this study. Blood was collected at baseline (POD0) and then on postoperative day (POD) 1, 3 and 28. Peripheral Blood Mononuclear Cells (PBMC) were isolated immediately and cryopreserved for flow cytometry and functional assays, including T and B cell ELISpot for measurement of IFNγ and IgG respectively, in response to the SARS-CoV2 spike (S) protein (RBD domain)or a non-specific stimulant (α-CD3 or α0IgG capture antibody, MT91 for T and B cells respectively) . Flow cytometry was used to characterize the cell surface phenotype of CD8+ T cells and intracellular IFNγ and TNFα. Serum was collected for quantification of antibody titres against the S-protein. Statistical comparisons were conducted on SAS® using the paired T-test and Wilcoxon Rank Test for parametric and non-parametric data respectively.

## Results

A total of 30 patients were included in the study. Surgical stress resulted in a significant 2-fold reduction in the proportion of CD8^+^ T cells that secreted IFNγ in response to the SARS-CoV-2 spike protein on POD1 by ELISpot (90.9±56.8 spots) as compared to POD0 (181.3±72.2, p<0.0001), which persisted on POD3 (128.5±49.2, p<0.01), but recovered by POD28 (163.2±51.9, p=0.1) (Figure 1). This was associated with a significant decrease in intracellular IFNγ and TNFα expression (p<0.05) and an increase in cell surface expression of the exhaustion marker, PD-1 (p<0.05). There was no change in the expression of exhaustion markers, TIM-3 or TIGIT, nor of T-bet, a marker of T cell effector function (Figure 2). Surgery also significantly impaired the production of IgG antibody by CD19^+^ memory B cells upon stimulation with the SARS-CoV-2 spike protein on POD1 by half (49.1±47.9 vs. 105.7±67.7 spots on POD0, p<0.0001). There was a measurable decrease in the IgM serum antibody titres against the S-protein on POD1 but not in the IgG or IgA serum antibody (Figure 2). In response to the non-specific stimulation, T cells and B cells were similarly highly dysfunctional as measured by IFNγ and IgG ELISpots respectively (Figure 1).

**Figure 1.**
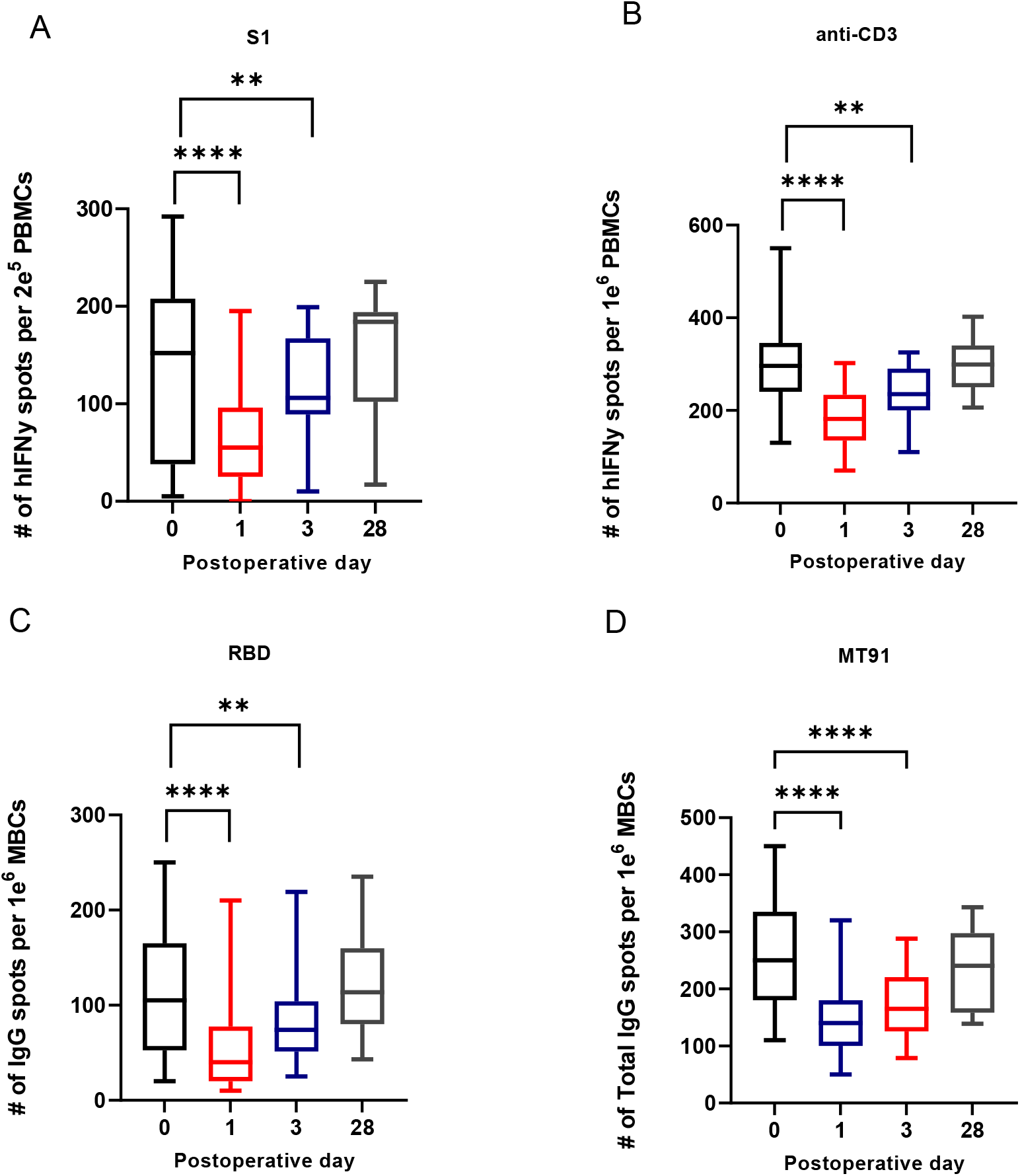
Surgical stress results in decrease in CD8^+^ T cell and memory B cell cytokine and antibody production in response to SARS-CoV-2 spike protein and RBD in vaccinated patients Adaptive immune responses by ELISpot against SARS-CoV-2 protein pool at pre-operative day 0 and post operative day 1, 3 and 28 in Pfizer vaccinated cancer patients (n=30). Proportion of CD3 + T cells producing hIFNγ against the spike S1 protein of SARS-CoV-2 peptide pool (A) anti-CD3 as positive control (B). Humoral immune responses by ELISpot in serum of vaccinated cancer patients. IgG responses to RBD of SARS-Cov-2 spike protein (C) total IgG response to MT91 stimulation (D) Data is presented as box plot with horizontal line in the middle representing the median. Statistical differences were analyzed by the paired T-test and Wilcoxon Rank Test. P values < 0.05 were considered significant.

**Figure 2.**
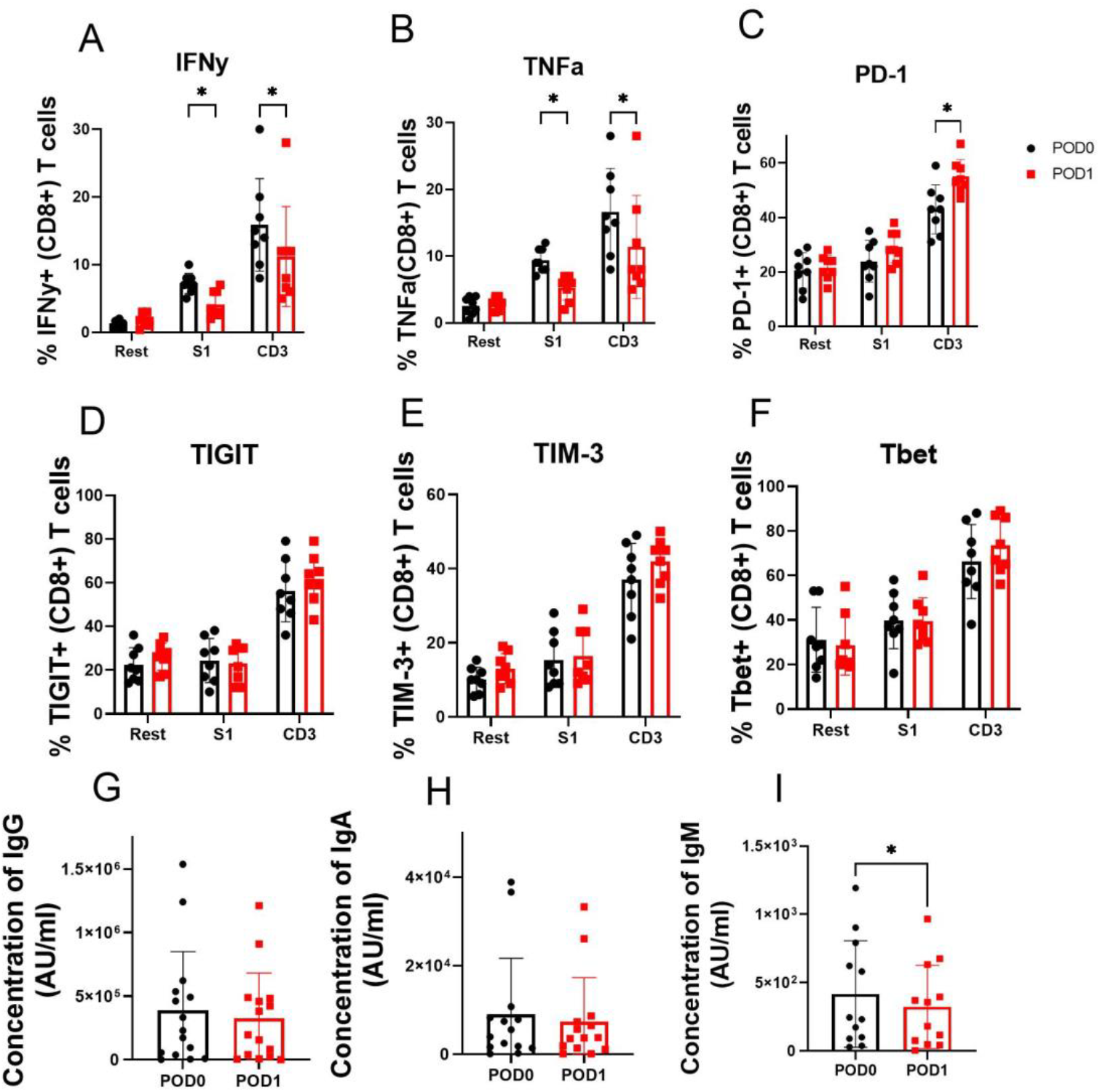
Effect of surgical stress on the regulation of IFN_γ_, TNFα, PD-1, TIGIT, TIM-3, T-bet in CD8^+^ T cells and characterization of serum IgG, IgM, and IgA in vaccinated patients CD8+ T cells from patients with profound suppression in adaptive immune response post-operatively from ELISpot assay (n=8), were further analysed by flow cytometry for changes in intracellular cytokine markers IFNγ (A), TNFα (B), exhaustion markers PD-1 (C), TIGIT (D), TIM-3 (E) and transcription factor Tbet (F). IgG (G), IgA (H) and IgM antibody levels were measured in serum of patients at pre-operative day 0 and post-operative day 1. Statistical differences were analyzed by the paired T-test and Wilcoxon Rank Test. P values < 0.05 were considered significant.

## Discussion

In the present study, we demonstrated the suppression of antigen specific CD8+ T-cell and B cell immunity following surgery in vaccinated cancer patients with a 2-fold reduction in immune response in the early postoperative period, using the SARS-CoV-2 S antigen. Given that patients were usually discharged by POD4, we were unable to clarify the duration of this dysfunction, but it has recovered by the end of the first postoperative month. The dysfunction of both important components of the adaptive and humoral immune system of cancer patients undergoing surgery may expose patients to not only an increased risk of cancer recurrence but also postoperative infectious complications. There are numerous studies correlating postoperative infectious complications^5^ and cancer recurrence and, more recently, studies demonstrating that minimal residual disease, as measured by circulating tumor (ct)-DNA at 1-month post-surgery, predicts cancer recurrence^6^. The obvious link between the two is postoperative immune suppression, which our lab has clearly linked in murine studies. This is the first study to clearly demonstrate profound antigen-specific T and B-cell dysfunction in the early postoperative period. Future research should focus on the mechanism of surgery-induced immune dysfunction which may aid in the development of therapeutics aimed at restoring it to improve both infectious and cancer-specific outcomes.

## Notes

### Competing Interest Statement

The authors have declared no competing interest.

